# Clumps: A Sequential Clustering Approach to Partitioning Sets of Phylogenies with Non-Identical Leaf Sets

**DOI:** 10.1101/2025.03.18.641816

**Authors:** Ana Serra Silva, Mark Wilkinson

## Abstract

Post-processing of sets of inferred phylogenetic trees often focuses on the canonical case of consensus of (multi)sets of phylogenetic trees on the same leaf set. However, with growing numbers of phylogenomic studies resorting to summary and/or supertree methods to obtain a phylogeny, the amount of (multi)sets of trees with non-identical leaf sets also increases. In an attempt identify trees with non-identical leaf sets that are topologically similar, we define a new sequential subsetting approach, “clumps of trees”, based on the distance between any tree in a set and the set’s supertree. While clumps were developed with trees with non-identical leaf sets in mind, they can be applied to (multi)sets of trees with identical taxonomic sampling. Unlike islands of trees, clumps will not always be mutually exclusive, thus making them more similar to families of trees.

---

Confronted with (multi)sets of inferred phylogenetic trees, common practice in systematic biology has been to represent them with a single consensus or supertree (Felsenstein, 2004). In contrast, several researchers have suggested that when sets of trees are not homogeneous it is better to partition the trees and summarise the subsets separately (e.g., Hendy et al., 1988; Maddison, 1991; Serra Silva and Wilkinson, 2021; Sumrall et al., 2001). In practice, most methods for discovering heterogeneity in and partitioning sets of trees have been developed for the special case where the trees have identical leaf sets, the so called ‘consensus case’ (e.g., Bonnard et al., 2006; Serra Silva and Wilkinson, 2021; Stockham et al., 2002). However, sets of trees on non-identical leaf sets are increasingly common, e.g. from genome-scale data. Thus, there is a need for methods for interrogating (multi)sets of trees in this, more general, ‘supertree case’ and partitioning them when they are found to be heterogeneous.

Sets of inferred trees may be heterogeneous due to real differences in evolutionary histories, such as those produced by gene duplication and loss, incomplete lineage sorting, etc. (e.g., Chan et al., 2020; Hahn, 2007), or because of incorrect inferences, such as those produced by systematic biases and/or stochastic and other errors (e.g., Ĺeveilĺe-Bourret et al., 2017; Simmons et al., 2022). Though we will not explore these, many important approaches to investigating heterogeneity focus on the underlying data from which trees are inferred (e.g., statistical binning, Bayzid et al. (2015); ortholog enrichment, Siu-Ting et al. (2019)). Other approaches that focus solely on the inferred trees have also been developed, almost exclusively within the consensus context (e.g., islands of trees, Maddison (1991) and Serra Silva and Wilkinson (2021); families of trees, Hendy et al. (1988); *k*-medoids clustering, Tahiri et al. (2018); multipolar consensus, Bonnard et al. (2006); consensus networks, Holland and Moulton (2003); tree alignment graphs, Smith et al. (2013)). These can be broadly divided into those based on split compatibility (e.g., multipolar consensus and tree alignment graphs) or on tree-to-tree distances, which seek to produce either consensus networks or to partition (multi)sets of trees into more homogeneous subsets (e.g., families and islands of trees, *k*-medoids clustering).

Here, we present the theoretical background for a novel tree-to-(super)tree distance-based approach to partitioning any heterogeneous (multi)set of trees, clumps of trees, that does not make use of data alignments and which can be applied in cases where the trees in a (multi)set have non-identical leaf sets. We further demonstrate the usefulness of clumps of trees by applying the *clumpy* pipeline to a series of empirical phylogenomic datasets.

## Defining Clumps of Trees

When dealing with (multi)sets of trees with the same leaf set it may be possible to identify mutually exclusive and exhaustive subsets that correspond to alternative evolutionary histories and/or analytical artefacts, which cause topological incongruence (e.g., Hendy et al., 1988; Maddison, 1991; Serra Silva and Wilkinson, 2021). However, identifying those subsets in (multi)sets of trees with non-identical leaf sets requires overcoming a number of obstacles, the most fundamental of which is the generalisation of tree-to-tree distances from the consensus to the supertree case.

This generalisation revolves, primarily, around how to make two distinct leaf sets identical, which can be achieved by pruning or grafting leaves to one or both trees, so as to render their leaf sets identical. In pruning, which we will be our framework hereafter, we use the intersection of the leaf sets to identify the subtrees induced by the shared taxa, whereas with grafting, we seek the pairs of (most similar) trees that display the original trees and the union of the leaf set (Cotton and Wilkinson, 2007). However, both approaches have difficulty dealing with trees with no or minimal overlap, which is particularly problematic for partitioning methods that require exhaustive pairwise distance matrices, e.g. islands of trees (Maddison, 1991; Serra Silva and Wilkinson, 2021). Islands are the disconnected components of a graph where vertices correspond to trees and edges connect all trees within a fixed distance threshold, and trees within islands may be quite dissimilar provided they are ‘connected’ by a series of sufficiently similar intermediates (Maddison, 1991; Serra Silva and Wilkinson, 2021).

Two imperfect solutions to the low leaf set overlap problem are to either set the distance between non-overlapping trees to zero, which can represent absence of conflict (e.g., Robinson-Foulds (RF, Robinson and Foulds, 1981) and quartet (QD, Estabrook et al., 1985) distances), or to treat these instances as non-applicable, which are generally dealt with by removal or replacement with imputed values (reviewed in Wagstaff, 2004). Neither solution is well suited for exhaustive pairwise distance methods like islands. Furthermore, even when all distances are defined, the search for islands, or any such clustering that relies on ‘connectedness’, can be derailed by insufficient overlap between trees because small trees can be sufficiently similar to larger, highly disparate trees, thus placing the larger trees in the same island and masking tree heterogeneity (Fig.1). This small tree problem might be alleviated by filtering trees under a minimum leaf set size. However, it still does not deal with pairs of non-overlapping trees, meaning that the question of how to define tree-to-tree distance(s) between non-overlapping trees remains.

**Fig. 1.**
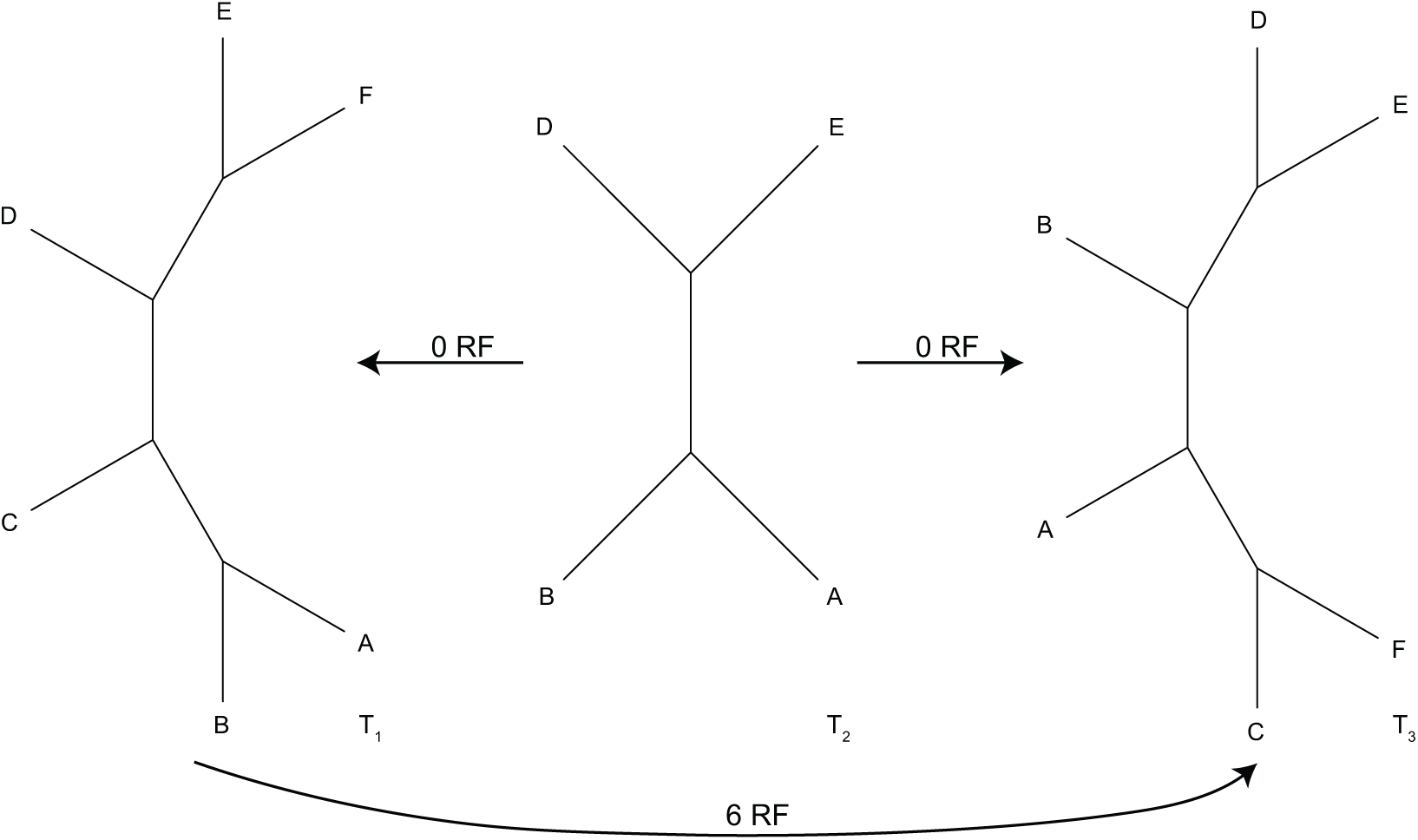
Small tree problem. Three partially overlapping trees where the two larger trees (*T*_1_ and *T*_3_) are maximally dissimilar under the RF distance, yet equidistant from *T*_2_. In this scenario, despite having no splits in common *T*_1_ and *T*_3_ would be placed in the same RF-island.

The issue of how to compare these minimally overlapping trees can be addressed by using a clustering approach that relies on the distance from a focal tree to all trees in the (multi)set. One such strategy is the use of families of trees (Hendy et al., 1988), where members of a subset are identified based on their distance to a tree *T*. If a (multi)set’s supertree were set as *T*, it would ensure that, because all trees in the set partially overlap with the supertree, there would be no undefined tree-to-supertree distances. Unfortunately, families of trees lack an unambiguous formal definition, making it unclear not only how tree *T* is selected, but also how clustering proceeds past the identification of the first family. To prevent potential ambiguities while attempting to extend families of trees to sets of trees with non-identical leaf sets, we define a new clustering approach called clumps of trees (a play on clump *n.*: 1.a. a compact mass, 2.a. a cluster of trees and clump *v.*: 2.b. to make into a clump, Oxford English Dictionary (OED)). We define clumps of trees as the sets that are sequentially extracted from a tree distribution, with each subset containing all trees in a distribution within a selected distance of the input set’s supertree, and any trees placed in a clump being removed from the partitionable distribution (Appendix I and Fig. 2). In set theory terminology, the partitionable tree distribution is the complement of the subset trees (S*^C^* in the formal definition below), i.e. the trees not yet assigned to a clump.

Formally, given i) a set Ƭ of trees and its supertree Ƭ*_ST_*, ii) a pairwise distance function 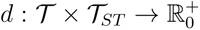, and iii) a threshold 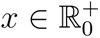, we define a sequence of sets S, such that 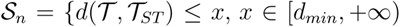 and Ƭ = S*^C^*}. The *tree clumps* of (Ƭ, *d*, *x*) correspond to the exhaustive subsets of the sequence of sets S. This definition allows for the threshold to be changed at each clump identification iteration, meaning that individual clumps can be defined on a different threshold distance to the supertree of the previous clump(s)’s S*^C^*. This flexibility in setting the threshold means that no trees are ‘unclumpable’, i.e. all trees will be placed in a cluster. We propose that the threshold distance be set at the troughs, points at which the sign of the first derivative of the density curve goes from negative to positive. Using histogram bin heights as proxy for the density curve, this allows for two ‘strict’ scenarios, one where every trough defines a clump (strict trough), the other where a clump is defined only when *y* = 0 (breaks), and a ‘loose’ scenario where the difference between the height of the nearest peaks and troughs (Δamplitude) is used as a guide of whether to partition the distance distribution (Fig.3). As in Serra Silva and Wilkinson’s (2021) generalised tree island definition, we do not specify a tree-to-(super)tree distance on which to define clumps. As such, any tree-to-tree distance that can be applied to distributions of trees with non-identical leaf sets can be used, and the choice of whether to work with (multi)sets and/or branch lengths is left to the user.

**Fig. 2.**
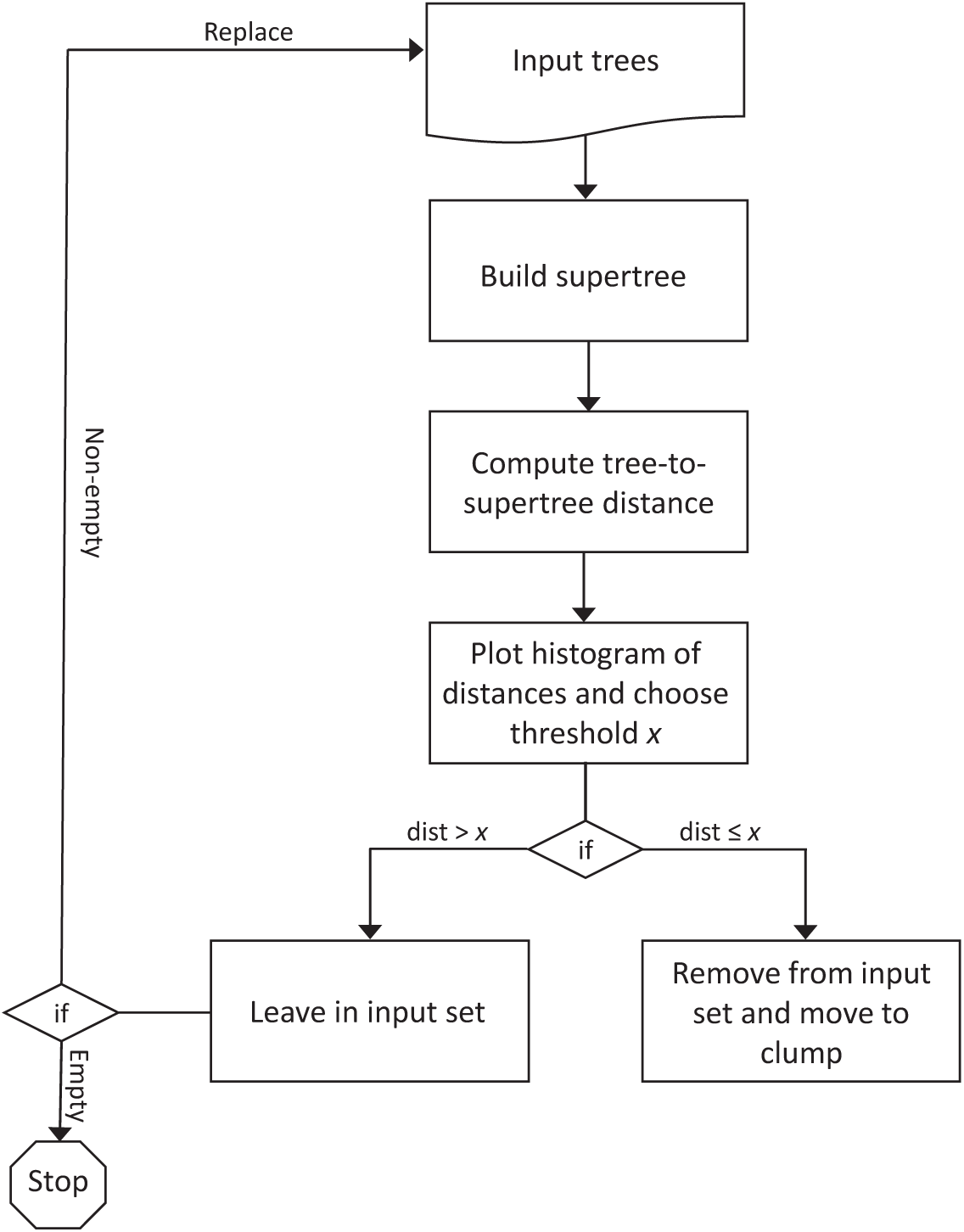
Clumping pipeline. Graphical depiction of the algorithm in Appendix I.

Three special cases follow easily from the definition of clumps of trees: i) at least one supertree corresponds to an input tree, ii) all supertrees correspond to input trees, and iii) if all input trees have the same leaf set, the supertree will also be a consensus tree. While it is not immediately apparent how these special cases might affect clump identification, the choice of supertree (or consensus) method will indubitably affect the identification of clumps, since the use of distinct supertrees to identify clumps from the same tree distribution may yield different tree clump numbers and/or structure.

Additionally, because clumps are defined on the distance to a single tree, like families of trees, the clumps may not be mutually exclusive. In other words, the same input tree might be within the desired distance threshold to multiple input supertrees, but once it has been assigned to a clump that input tree is no longer available for comparison to other supertrees. While under identical analytical settings the same trees will be placed into the same clumps, changing bin sizes may lead to slightly different clump structures being identified, and not necessarily by the process of clumps merging (as is seen when island extraction thresholds are increased, see Serra Silva and Wilkinson (2021)), see below. This will be of particular importance when dealing with sets of trees with non-identical leaf sets, since, if using uncorrected distances, the smaller subtrees will fit in multiple clusters despite their expected placement within the first clump(s) to be extracted. However, as stated above, trees with fewer than a selected number of leaves can be filtered from the initial tree distribution, thus reducing the number of trees that might fit into multiple clumps.

**Fig. 3.**
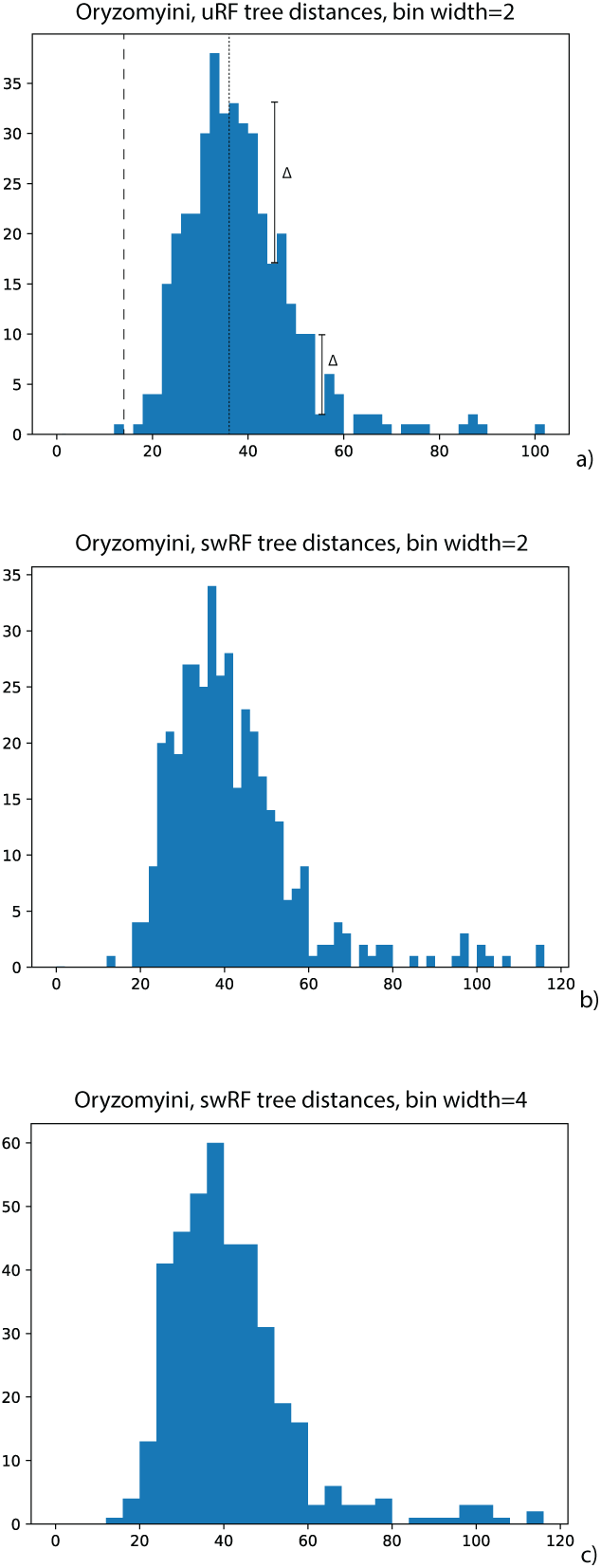
Example of the histograms produced during the clump extraction pipeline (see Appendix I and Fig. 2), using Percequillo et al.’s (2021) Oryzomini dataset, under a) uncorrected Robinson-Foulds distance (uRF, Robinson and Foulds, 1981) and bin width of two, b) size-weighted RF distance (swRF) and bin width of two, and c) swRF and bin width of four. In a) the dashed line illustrates the break (y=0) partitioning approach, the dotted line illustrates the strict trough approach (after removal of the clump at RF=14), and the line segments illustrate the user-defined Δamplitude approach.

## Leaf Set Size-Aware Tree-to-Tree Distance

A final technical consideration before demonstrating the use of the *clumpy* pipeline is our choice of tree-to-supertree distance. As noted above, clumps can be defined on any tree-to-tree distance. However, because we are dealing with trees with non-identical leaf sets care is required when interpreting and/or using uncorrected tree-to-supertree distances, since they actually correspond to the distance between each input tree and the subtree they induce on the supertree (Cotton and Wilkinson, 2007). This leads to situations where the same RF distance between two trees, of different sizes, and the supertree may convey different levels of topological divergence (Fig.1). For example, assuming binary trees, a maximally divergent tree-supertree pair with only four shared tips will have an RF of 2, that same RF in any tree-supertree pair with over four tips in common conveys the minimal non-zero topological divergence between trees. This means that the differences in size between input trees and the supertree are not taken into account in uncorrected tree-to-supertree distances.

Using the RF distance, due to its ease of implementation and interpretation, we propose a modification of the uncorrected RF distance (uRF) that accounts for the theoretical maximum RF for the supertree, we are calling this modification the size-weighted RF distance (swRF). The swRF is computed using

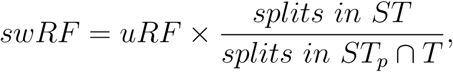

where “splits in *ST_p_*∩ *T*” is the intersection of splits in the pruned supertree *ST* and in tree *T*. This calculation is implemented in the supertree-based clump extraction pipelines (see Appendix I and Fig. 2) with ETE’s v.3.1.1 (Huerta-Cepas et al., 2010) RF calculator, where the uRF is calculated directly and the swRF calculation uses the edge inclusion/exclusion information outputted by the RF calculator. However, other modifications to the RF distance, specifically for trees on non-identical leaf sets, are available (e.g., Llabŕes et al., 2021; Tahiri et al., 2022).

**Table 1:** Molecular datasets used for clumping analyses. Datasets starting with *KST* refer to Siu-Ting et al. (2019). In the datatype column, AHE corresponds to anchored hybrid enrichment, and UCE to ultraconserved elements.

| Dataset | Ingroup taxa | Datatype | Trees | Taxon sampling | Supertree size |
| --- | --- | --- | --- | --- | --- |
| Percequillo et al. (2021) | Oryzomyini | AHE | 402 | 38–75 | 75 |
| Hime et al. (2021) | Lissamphibia | AHE | 220 | 201–290 | 291 |
| Streicher et al. (2020) | Microhylidae | UCE | 1796 | 24–49 | 64 |
| KST2656 | Lissamphibia | Transcriptome | 2656 | 4–32 | 33 |
| KST2019 | Lissamphibia | Transcriptome | 2019 | 4–32 | 33 |
| KST768 | Lissamphibia | Transcriptome | 768 | 6–32 | 33 |
| KST348 | Lissamphibia | Transcriptome | 348 | 6–32 | 33 |

## Examples

To test multiple aspects of clumping we will be using five distinct tree sets from studies that represent commonly used types of phylogenetic data, including discrete morphological characters, and molecular sequences obtained using anchored hybrid enrichment (AHE), ultraconserved element (UCE) enrichment and transcriptomics (table 1).

First, we will use Percequillo et al.’s (2021) AHE-based dataset to illustrate that clumping analyses can and do identify differing evolutionary histories present in the tree set. Given the supertree/supermatrix conflict they reported for oryzomyine rodents’ clade D (Fig. 1 in Percequillo et al., 2021), we expect the clumping analysis to identify at least two major topological variants in two clumps. Since all trees in the set have at least 50% of the sampled taxa present, the effects of tree size and complete non-overlap between trees on clumping cannot be addressed with this dataset.

We will be using a second AHE dataset (Hime et al., 2021), primarily, to test whether changing histogram bin width affects clump identification, for details on the analytical pipeline see Appendix I and figure 2. While all datasets were clumped under multiple bin widths, the Hime et al. (2021) dataset was chosen specifically to test this analytical parameter for two reasons. First, the dataset was originally used to explore deep-time relationships between Lissamphibia (the least inclusive clade containing Gymnophiona, Caudata and Anura), and was shown to contain at least three major topological variants, corresponding to distinct lissamphibian evolutionary histories: i) Batrachia (Caudata+Anura), ii) Procera (Caudata+Gymnophiona), and iii) Acauda (Anura+Gymnophiona). Secondly, of the selected datasets, Hime et al.’s (2021) has the largest trees (table 1) and, thus, the widest range of possible RF distances between trees (*RF* ∈ [0, 576]). As such, by modifying the bin width parameter we can not only compare how the number of clumps changes between analyses, but also which main topological variants are identified, and how the proportion of clumps supporting each hypothesis changes between analyses.

Another type of anchored markers commonly used in phylogenetic analyses are UCEs, which differ from AHEs by having shorter, less divergent sequences (Singhal et al., 2017). Because of UCEs’ popularity we analysed a dataset previously used to attempt to resolve recalcitrant branches in Microhylidae (Streicher et al., 2020). Because the largest tree(s) in the microhylid dataset do not exceed 77% taxon sampling (the lowest observed maximum taxon sampling of all datasets), this dataset allows us to explore how much ineffective overlap affects clumping analyses.

The final dataset based on molecular data we will test is Siu-Ting et al.’s (2019) transcriptome-based tree (sub)sets, which were originally used to explore the effects of paralog inclusion on topological incongruence, branch support and divergence estimates on lissamphibian relationships. This dataset, KST hereafter, is particularly interesting because it is made up of multiple subsets of a larger tree distribution, which allows for the exploration of whether the number of trees in a set affects clump identification, and might shed light on whether the topologies most similar to a dataset’s summary tree are found in the largest clump(s), the most clumps, or a combination of both. And, given that two of the subsets include quartet trees, we can test the assumption that under uncorrected distances the smallest trees are indeed swept into the first clump(s).

Lastly, the set of most parsimonious trees (MPTs) recovered from the morphological data matrix used by Pardo et al. (2017) to place *Chinlestegophis jenkinsi* in a phylogeny of fossil amphibians will be used to explore some of the special cases identified above. Particularly, how changing supertree inference method might influence the clumps that are identified. As one of the tree distributions used to illustrate Serra Silva and Wilkinson’s (2021) generalised definition of islands of trees, we will also compare the output of the clumping analyses, under the strict trough and break scenarios, to the known 10- and 12-RF island sets. We will also use this tree distribution to compare our clustering approach to Tahiri et al.’s (2022) *k*-means clustering algorithm for trees with partially overlapping leaf sets.

## Extracting Clumps from Distributions of Trees with Non-Identical Leaf Sets

Before delving into the biological patterns identified by the clumping of each dataset, it is worth noting that, for the majority of the analyses, as the number of extracted clumps increases so does the dissimilarity between the trees within clumps and the supertree of all input trees (ST_all_), with many of the clump supertrees displaying complete breakdown of ingroup monophyly (from species to class). While results for the uRF analyses will not be shown, they confirmed our conjecture that, under an uncorrected tree-to-supertree distance, all quartet trees are placed into the first clump (not the case with the swRF). We did not, however, explore up to what size tree this remains true. A breakdown of the number of clumps extracted from each tree distribution, under swRF and multiple bin widths, can be found in table 2 and all the files resulting from the analyses are available in the Dryad Repository https://dx.doi.org/TBD. Lastly, while the pipelines were designed to allow the users to choose the threshold, and thus which of the suggested approaches to clump identification to use (Fig.3a), unless stated otherwise, all results below correspond to the strict trough approach with bin width of two.

Starting with the rodent dataset, before clump supertrees start displaying extensive non-monophyly (*c.* clump 14) the differences in topology between the clumps are mostly restricted to the taxa in clade D (see figure 1 in Percequillo et al. (2021)), particularly *Cerradomys*, *Lundomys* and *Sooretamys*, which are also the taxa responsible for the supertree/supermatrix incongruence reported by Percequillo et al. (2021). While clades A and B display some instability, it is the taxa in clade C that are the most unstable after those in clade D, with at least three clump supertrees displaying C paraphyletic with D. From a clump identification standpoint, this is an interesting dataset since the first clump is composed of a single tree that is not topologically identical to ST_all_, yet the supertree of clump-2 is identical to ST_all_. Moreover, this pattern was found both with the uRF and swRF analyses. Thus, while the first clump contains those trees that are the most similar to ST_all_, the clump summary that best matches ST_all_ may not correspond to that of the first clump. When bin width is increased, very little information can be gleaned from the outputted clumps. With a bin width of four, two clumps are still recovered that show clade D’s instability in comparison to the other oryzomyine clades. But, at higher bin width values, apart from a single clump whose supertree matches ST_all_, only clumps exhibiting a combination of rooting and non-monophyly of all/most Oryzomyini clades are identified. As such, for this dataset the most informative bin width setting is bin=2.

**Table 2:** Number of clumps identified for each dataset under the strict trough approach. The *bin#* column names refer to the histogram bin width used for each analysis.

| Dataset | bin2 | bin4 | bin6 | bin8 | bin10 | bin12 | bin14 |
| --- | --- | --- | --- | --- | --- | --- | --- |
| Percequillo et al. (2021) | 26 | 12 | 9 | 7 | 7 | 5 | 1 |
| Hime et al. (2021) | 41 | 21 | 15 | 13 | 6 | 6 | 11 |
| Streicher et al. (2020) | 20 | 3 | 1 | 1 | 1 | 1 | 1 |
| KST2656 | 38 | 5 | 2 | 2 | 1 | 1 | 1 |
| KST2019 | 52 | 15 | 4 | 2 | 1 | 1 | 1 |
| KST768 | 24 | 2 | 2 | 1 | 1 | 1 | 1 |
| KST348 | 22 | 7 | 1 | 1 | 1 | 1 | 1 |

Unlike the previous dataset, none of the analyses on the Hime et al. (2021) data yielded a clump with its supertree identical to ST_all_. For the default analysis (bin width=2), the summary trees most similar to ST_all_ correspond to those of clumps five and eight, yet their distance to the supertree is of 40 RF. In the analysis with bin width of 4, the minimum distance between the supertrees of the clumps and ST_all_ is 12 RF and corresponds to clump-2. The analyses with the lowest distance between a clump supertree and ST_all_ are those under bin widths 12 and 14, where the supertree of the first clump is 4 RF away from ST_all_, with the difference between the supertrees revolving around the relationship between Nyctibatrachidae and Ceratobatrachidae. Another major difference between the analyses with bin widths two and four is the nearly halved number of identified clumps when the histogram bin width is doubled, see table 2. Interestingly, when the clumps with outgroup rooting, extensive ingroup non-monophyly and taxon sampling issues are ignored the default analysis yields a breakdown of clumps supporting Batrachia, Procera or Acauda similar to that reported by Hime et al. (2021), though the proportions are not an exact match to Hime et al.’s 2021 analysis. They identified a proportion of trees supporting Batrachia:Procera:Acauda of approximately 2:1:1, and the default clumping analysis yielded a proportion of 3:1:1. We should note that this discrepancy in the proportions might be partly due to the fact that we kept all gene trees for the clumping analyses, while Hime et al. (2021) filtered out all trees that could not be used to test the relationship between lissamphibian orders. In the bin width=4 analysis, ignoring clumps with Lissamphibia non-monophyly issues, no clumps support Acauda, a single clump supports Procera and nine Batrachia. In both the bin=2 and bin=4 analyses, Gymnophiona is the most internally stable amphibian order, the majority of trees that include Sirenidae and Cryptobranchoidea have the latter as sister to all other caudates, and most of the intraorder taxonomic instability is found within Anura, specifically Neobatrachia. At larger bin widths, the initial clumps all recover Batrachia, but any clump supporting Procera or Acauda has Amniota nested within Lissamphibia and *Latimeria* as sister to it. Unlike the rodent dataset, the non-monophyly of the ingroup (Lissamphibia) is mostly due to paraphyly with the outgroups, not to para- and polyphyly of the ingroup’s lower taxa.

The microhylid UCE dataset (sampling of 37.5%–76.5%) yielded 20 clumps, under the bin width=2 analysis, with RF=28 being the minimum distance between a clump supertree, with all 64 taxa, and ST_all_. Most clump supertrees exhibit extensive non-monophyly at the subfamily level, which is likely the result of ineffective overlap within clumps, this is congruent with the large number of low support branches in the ST_all_ (figure 2.B in Streicher et al., 2020) and the supertree/supermatrix incongruence reported in the original paper. Given that no tree in this dataset exceeded a taxon sampling of 77%, the lowest maximum observed sampling of all analysed datasets, it is not unreasonable to think that this is the parameter causing the clumping pipeline’s inability to identify ‘major’ topologies. The uRF analysis of this dataset yielded similar results, but found only 9 clumps and, more importantly, a considerably lower proportion of unresolved clump summary trees (swRF=0.8, uRF≈0.4). The latter suggests that poor effective overlap within clumps is indeed the culprit of the patterns identified in the swRF analysis, since the numbers of unresolved clump supertrees greatly decreased when clump size increased (uRF). This is further cemented by the recovery of three large clumps (*>* 300 trees) whose summaries display primarily monophyletic microhylid subfamilies when the bin width is increased to four and a minimal distance to ST_all_ of 24 RF, although any further increase in bin width leads to the recovery of a single clump.

The KST datasets broadly follow the patterns reported by Siu-Ting et al. (2019) for their supertree analyses, with the majority of clump supertrees (that do not have extensive outgroup rooting, non-monophyly and/or taxon sampling issues) supporting the same hypothesis of amphibian relationships as the ST_all_ of each dataset. In other words, most clump supertrees from KST348 and KST2019 support Batrachia, and those for KST768 and KST2656 support Procera. Two things of note that can be found in all analyses of the KST datasets are that Anura is the most internally unstable amphibian clade, and that the supertree of clump-2 is either identical or the most similar to ST_all_. For KST2019 and KST2656, the latter observation can be explained by 50% of the first clump consisting of quartet trees, which leads to increased topological instability. For KST348 and KST768, the reason is not as clear, though that the proportion of small trees (6-tip) is higher for clump-1 than for clump-2 may contribute to the lower resolution and topological changes compared to ST_all_. As for the non-default bin width analyses, they revealed a pattern similar to Hime et al.’s (2021), but from a bin width of 10 all datasets recovered a single clump (table 2).

Overall, the results show that clumping can extract some of the major topologies found in (multi)sets of trees, but the effects of a set’s maximum tree size on the analyses need to be explored further. Perhaps more importantly, clump informativeness can be manipulated by varying the histogram bin width, with some datasets benefiting from wider bins, like the microhylid trees, whereas others benefit from narrow bins, much like the Oryzomyini tree set.

## Special Cases

In the section describing clumps of trees, we introduced a number of special cases that are expected to occur in clumping analyses. The Pardo et al. (2017) MPTs allow us to explore two of those, the case where most supertrees correspond to input trees and the use of consensus trees. The latter also helps to illustrate how changing the supertree method may change the clumps identified. Additionally, because the tree-to-supertree distance histogram for all MPTs shows a distribution made of two connected peaks and a third disconnected peak, this dataset is ideal to illustrate the differences between using the strict trough and break approaches to identify clumps. Under the break approach the *clumpy* pipeline that computes the supertrees using Astral-III v.5.7.7 (Zhang et al., 2018) identifies four clumps that correspond to the four 12-RF islands reported in Serra Silva and Wilkinson (2021). The trough analysis, however, identified the three smallest 10-RF islands as clumps, but the two largest clumps do not map onto the two largest 10-RF islands. Thus, while providing information similar to that of islands, regarding major hypotheses for evolutionary histories, clumps of trees on the same leaf set are not always identical to tree islands. However, with large tree distributions clump identification is less memory intensive than island extraction analyses and allows for exploration of topological heterogeneity, even when RF-island identification fails (e.g., Serra Silva, 2024).

## Using a majority-rule consensus as the supertree

Using the majority-rule consensus (MRC, Margush and McMorris, 1981) as the supertree, computed with DendroPy v.4.5.2 (Sukumaran and Holder, 2010), one immediate change between the MRC and Astral results is the right-displacement of the distance distributions. This is due to the MRC not being one of the input trees, unlike most of the summary trees computed by Astral, thus increasing the minimum distance between the ‘supertree’ and all input trees. As for what clumps are identified, under the break approach the four 12-RF islands are recovered once more, though the distance thresholds differ between the Astral and MRC analyses. The trough analysis yields five clumps, with the three smallest corresponding to the smallest 10-RF islands, but the two largest clumps not matching the two largest 10-RF islands, much like in the supertree case above.

Additionally, the make-up of the two largest clumps is distinct from those obtained in the Astral+trough analysis. Thus, by changing the supertree method we can change the number and size of the clumps that are identified from the same dataset.

**Table 3:** Taxon rarefaction analyses. Number of clumps identified under the swRF and a bin-width of 2. . Negative numbers in header correspond to the number of taxa removed for each analysis.

| Replicate | -2 taxa | -4 taxa | -8 taxa | -16 taxa | -32 taxa |
| --- | --- | --- | --- | --- | --- |
| 1 | 4 | 4 | 5 | 14 | 47 |
| 2 | 4 | 4 | 5 | 19 | 42 |
| 3 | 4 | 4 | 5 | 21 | 35 |
| 4 | 4 | 4 | 5 | 14 | 40 |
| 5 | 4 | 4 | 5 | 17 | 36 |
| 6 | 4 | 4 | 7 | 17 | 45 |
| 7 | 4 | 4 | 5 | 16 | 35 |
| 8 | 4 | 4 | 7 | 14 | 45 |
| 9 | 4 | 4 | 5 | 16 | 42 |
| 10 | 4 | 4 | 5 | 17 | 36 |

## Rarefying MPTs to explore the effects on ineffective overlap

Clumping analyses of the microhylid dataset suggested that there may be a minimum amount of maximum taxonomic sampling for all trees in a distribution for the pipeline to yield useful tree clusterings. Because we know the behaviour of Pardo et al.’s (2017) MPTs under multiple clumping and island thresholds/conditions, we set up a small experiment with taxonomically rarefied sets of this MPT distribution. The set of MPTs was subjected to the removal of 2, 4, 8, 16 or 32 random taxa, with 10 replicates each, for a maximum taxonomic sampling per tree of approximately 97%, 95%, 89%, 76% and 58%. The distributions of rarefied trees were run through the *clumpy* pipeline under the swRF with a bin width of 2, and the break approach for threshold selection (table 3).

As expected, the tree sets with two or four taxa removed yielded sets of clumps that are very similar (occasionally even identical) in number and make-up to those identified from the original MPTs, as well as the previously identified 12-RF islands. However, even from 11% missing taxa per tree the clump structure veers drastically from that identified from the original set of MPTs. At first glance it may appear that the clumps identified from most of the -8 taxa replicates match the 10-RF islands, but the size of the clumps does not match that of the islands. This means that ineffective overlap is already apparent, but not yet strong enough to obscure the presence of topological variants. By the time we are removing 16 taxa (very similar to the 77% taxon coverage of the microhylid trees), though, the breakdown in the informativeness of clumps starts becoming apparent with the recovery of 14–21 clumps per replicate (table 3). Increasing bin width to 4 brings the number of clumps down to four (table 4), confirming that bin width is an important aspect of the clumping analyses and that this parameter may smooth some of the noise introduced by lack of resolution due to poor taxonomic overlap.

**Table 4:** Number of clumps identified for the first replicate of each taxon rarefaction analysis under the strict break approach. The *bin#* column names refer to the histogram bin width used for each analysis.

| Dataset | bin2 | bin4 | bin6 | bin8 | bin10 | bin12 | bin14 |
| --- | --- | --- | --- | --- | --- | --- | --- |
| -2 taxa | 4 | 4 | 4 | 4 | 4 | 4 | 4 |
| -4 taxa | 4 | 4 | 4 | 4 | 4 | 4 | 4 |
| -8 taxa | 5 | 4 | 4 | 4 | 4 | 4 | 4 |
| -16 taxa | 14 | 4 | 4 | 4 | 4 | 4 | 4 |
| -32 taxa | 47 | 4 | 4 | 4 | 1 | 1 | 1 |

## Comparing clumping and *k*-means clustering

Recently, Tahiri et al. (2022) have developed a supertree clustering approach using *k*-means with similar aims to the clumping approach described here. As with our swRF, they also developed a ‘corrected’ RF calculation, but while ours accounts for the maximum possible RF given the ‘unpruned’ supertree, Tahiri et al.’s (2022) normalises the RF given the pruned supertree. However, their clustering analyses does not use that normalised distance directly, but an Euclidean approximation to the *k*-means objective function, with a penalisation parameter to prevent trees with little taxonomic overlap from being placed in the same cluster.

Using the first replicate generated by removing two random taxa from the set of Pardo et al.’s (2017) MPTs as input, we compared the output of the clumping and *k*-means approaches. Due to memory restrictions the tree file had to be reduced to 235 trees in order to successfully run the *k*-means clustering, under the recommended validity index (Calinski-Harabasz), a minimum of two clusters with a maximum of 234 trees, and the penalisation parameter set to zero (to approximate our swRF calculation). With these parameters the *k*-means analyses identified four clusters with 18, 72, 90 and 55 trees, identical to the clumps identified both under uRF and swRF. This shows that our clustering approach does identify topologically-relevant clusters of trees, and at this time is the only clustering software for trees with non-identical leaf sets that can handle large input files and single-cluster tree sets.

## Discussion

There is a long history of post-processing (multi)sets of phylogenetic trees to find the most informative summary for these sets by clustering and/or building consensus trees (e.g., Bonnard et al., 2006; Hendy et al., 1988; Maddison, 1991; Serra Silva and Wilkinson, 2021; Stockham et al., 2002). Many post-processing approaches focus on sets of trees with completely overlapping leaf sets, but with the increased use of phylogenomic datasets comes the need for tools that can be applied to sets of trees with non-identical leaf sets. First, since supertrees are an extension of consensus methods (Gordon, 1986), in the context of trees with non-identical leaf sets we may consider them to fulfil the same role as consensus methods do when dealing with sets of trees on the same leaf set. A non-supertree ‘consensus’ method that may at first appear enticing and would allow for the identification of a set of relationships/taxa present in all input trees is computing the maximum agreement subtree (MAST, Amir and Keselman, 1997; Finden and Gordon, 1985). However, as the number of input trees and/or polytomies increases, the complexity of computing the MAST for a set of trees becomes NP-hard (reviewed in Deepak et al., 2014), making this approach suboptimal when dealing with large tree distributions, which many phylogenomic datasets are. This makes clustering approaches the better option to post-process sets of trees with non-identical leaf sets.

However, clustering methods are not without challenges. While some of the main considerations for methods to cluster (multi)sets of trees on the same leaf set might entail rootedness and the presence of duplicate trees, for trees with non-identical leaf sets it is also necessary to take the presence of small trees (triplets/quartets), and of trees with little to no taxonomic overlap into account.

The presence of small trees is an important consideration seeing as two trees with highly disparate topologies, supporting distinct evolutionary histories (e.g., *T*_1_ supports Batrachia and *T*_2_ supports Acauda), might still display shared quartet/triplet trees. In island-like clustering methods, these quartets/triplets have the capacity to cluster highly disparate trees into the same subset, since the small distances between the quartet/triplet and the larger trees artificially lower the ‘connectedness’ threshold between the large trees (Fig.1). By implementing a sequential clustering approach, we need not worry about quartet trees affecting the clustering process, since, when using uncorrected distances, they were always placed in the first cluster(s) to be identified. However, further work is required to check up to what size tree we can expect to find the same pattern exhibited by the quartet trees, and whether size filtering prior to clustering changes the number of clumps identified. If using a corrected distance, like the swRF, quartet trees will be more widely distributed across clumps, provided more than one clump is identified, but each clump will only contain those quartet trees that perfectly match their input supertrees.

An interesting phenomenon that may occur with clumping is the identification of first clumps whose summary tree differs from the ST_all_, followed by a second clump whose summary tree is identical to the ST_all_. From the trough-based analyses we identified two distinct scenarios for the occurrence of this phenomenon. One scenario consisted of a small clump followed by a break and a large second clump, and the other scenario of a large first clump followed by a trough and a second large clump. The latter, and arguably more interesting, scenario was found in many of the KST datasets, where the summary tree of a large first clump was highly unresolved, yet the summary of the second clump matched the ST_all_. In this case, the first clump consisted of many (100s–1000s) trees, which had considerably fewer tips than the ST_all_ with poor effective overlap between them resulting in a highly unresolved summary tree. For both scenarios, it may be sensible to collapse the first two clumps into a single clump, and, more broadly, it may be sensible to collapse any clumps whose input supertrees are identical (superclumps).

Another possible cause of poor clump supertree resolution, not apparent from the tested tree sets, is the presence of subsets of trees equidistant to the input supertree, but not to each other. This would result in the clumping analysis being unable to separate the subsets and potentially obscure topological variants present in the tree set of interest. Currently, the only way to deal with this scenario would be to re-run *clumpy* calling a different supertree inference software, or combine the clumping analysis with another clustering approach (e.g., Tahiri et al., 2022).

With the exception of the microhylid dataset, under a bin width of two all analyses yielded a collection of clumps from which biological information can be gleaned. For example, from the KST and Hime et al. (2021) datasets it was possible to identify clumps that support the Batrachia, Procera or Acauda hypotheses of amphibian relationships. However, in all analyses a large number of clumps exhibits ingroup non-monophyly, which hinders the identification/interpretation of the summary trees’ and the biological information they display, whether it be at the order or genus level, and renders the clumps uninformative. This pitfall might be avoided either by filtering out those unrooted trees without ‘monophyletic’ outgroups, similar to the filtering used by Hime et al. (2021), or by using ingroup-only trees as input and rooting the clump summary trees with an *a posteriori* method (e.g., MAD, Tria et al., 2017).

The microhylid dataset presents its own set of pitfalls. While not unexpected, given the incongruence between the topologies recovered by the supermatrix and supertree analyses reported by Streicher et al. (2020), the level of dissimilarity between the summary tree of each clump and both the supertree of all trees and the supertree of each complement (S*^C^*) cannot be explained entirely by the presence of small trees, nor by outgroup polyphyly. Rather, that no tree in the set has a taxon sampling of over 77% suggests that there is extensive ineffective overlap between the trees in each clump. Especially, given that when bin width is increased, fewer and larger clumps are identified, with resolved and informative clump supertrees. This pattern was confirmed with the rarefied Pardo et al. (2017) tree sets, where the number of clumps identified under bin with of 2 increased substantially between the 89% and 76% maximal taxonomic coverage tree sets, and again yielded more similar numbers of clusters when bin width was increased (tables 3 and 4). Thus, while more testing will be required, it appears that, though clumping does not require datasets where at least some trees have 100% taxon sampling, it might have an optimal lower bound for the maximum observed taxon sampling, and that choice of bin width is an important analytical parameter. Additionally, the choice of supertree method may also be a consideration, since, as shown by the analyses on the Pardo et al. (2017) MPTs, different supertrees can yield different clump assortments. It is thus possible that if a different supertree method had been used, clump summaries might have imparted more information on alternate microhylid topologies.

Despite the myriad pitfalls described above, variable threshold clumping can and does yield relevant and useful information of the distinct evolutionary histories encoded in (multi)sets of trees. While the choices of supertree method, partitioning strategy and histogram bin width can influence the clumps identified, this new partitioning strategy has the benefit of not requiring data alignments nor phylograms (e.g., Smith et al., 2020), meaning that it can be applied to any (multi)set of trees. Additionally, by relying on a tree-to-supertree distance the tree order in the input file will not affect the analyses, unlike some of the currently available methods for post-processing trees with non-identical leaf sets (e.g., Smith et al., 2013). However, further work is required to identify the optimal minimum observed taxon sampling and outgroup monophyly parameters for the method.

While we used a modified RF distance for the analyses, there is a push to develop tree-to-tree distances specifically designed to deal with trees with non-identical leaf sets (e.g., leaf removal, Chauve et al. (2017); completion based RF, Bansal (2020)), which may either lead to other partitioning strategies or to a refinement of the definition of clumps based on the novel tree distance measures, which may in turn make the choice of partitioning scheme and/or histogram bin widths unnecessary. Recently, Tahiri et al. (2022) extended the single *vs*. multiple consensus debate to the supertree context with their proposed extension of *k*-means clustering to a pruning-based normalised RF distance. However, despite highly promising results, their method is unable to address the issue of distances between trees with two or fewer tips in common, and as shown above cannot handle large tree files. While *k*-means are less robust than other clustering approaches like *k*-medoids (Arora et al., 2016), Tahiri et al.’s (2022) work is a huge step in the post-processing of sets of trees with non-identical leaf sets, and may soon lead to a proliferation of generalised clustering approaches and other methods to rival the breadth of tools available for post-processing of trees on the same leaf set.

Lastly, with small alterations to the pipeline using the appropriate supertree and/or consensus software and a corresponding distance measure, the existing clump extracting pipelines can be extended to deal with (multi)sets of rooted trees and of internally labelled trees. The latter might consist of phylogenetic trees with branch labels corresponding to higher taxonomy, thus allowing for trees at distinct taxonomic levels to be compared (e.g., MultiLevelSupertree, Berry et al., 2012), or they might consist of the node-labelled mutation trees commonly used in cancer phylogenetics (e.g., Aguse et al., 2019).

## Acknowledgements

We thank Jeff Streicher, Benoit Morell, Tom Williams and James Cotton for helpful and thought provoking discussions about our clustering method. We also thank Miranda Sherlock for insightful comments of earlier versions of this manuscript. This work was supported by the Natural Environment Research Council [NE/L002434/1].

## Supplementary Material

Data available from the Dryad Digital Repository: http://dx.doi.org/TBD. *clumpy* Python pipeline available on GitHub: https://github.com/anaserrasilva/clumpy.

